# Laminar perfusion imaging with zoomed arterial spin labeling at 7 Tesla

**DOI:** 10.1101/2021.04.13.439689

**Authors:** Xingfeng Shao, Fanhua Guo, Qinyang Shou, Kai Wang, Kay Jann, Lirong Yan, Arthur W. Toga, Peng Zhang, Danny JJ Wang

**Affiliations:** Laboratory of FMRI Technology (LOFT), USC Stevens Neuroimaging and Informatics Institute, Keck School of Medicine, University of Southern California, Los Angeles, CA 90033, USA; State Key Laboratory of Brain and Cognitive Science, Beijing MRI Center for Brain Research, Institute of Biophysics, Chinese Academy of Sciences, Beijing 100101, China; Department of Neurology, Keck School of Medicine, University of Southern California, Los Angeles, CA 90033, USA; Laboratory of Neuroimaging, USC Stevens Neuroimaging and Informatics Institute, Keck School of Medicine, University of Southern California, Los Angeles, CA 90033, USA

**Author notes:** Correspondence: Danny JJ Wang. **Author Contributions**: X.S. implemented the MRI pulse sequence, conducted the experiments and data analysis. D.W., P.Z and A.T. provided advice on experimental design and research direction. X.S., D.W. and P.Z. designed the experiments. F.G., Q.S., K.W., L.Y. and K.J. provided input on the design of the analysis. All authors wrote the manuscript.

**Keywords:** Laminar fMRI, Perfusion, Arterial spin labeling, Neural circuit, Visual spatial attention, Ultrahigh field

## Abstract

Laminar fMRI based on BOLD and CBV contrast at ultrahigh magnetic fields has been applied for studying the dynamics of mesoscopic brain networks. However, the quantitative interpretations of BOLD/CBV fMRI results are confounded by different baseline physiology across cortical layers. Here we introduce a novel 3D zoomed pseudo-continuous arterial spin labeling technique at 7T that offers the unique capability for quantitative measurements of laminar cerebral blood flow (CBF) both at rest and during task activation with high spatial specificity and sensitivity. We found arterial transit time in superficial layers is ∼100 msec shorter than in middle/deep layers revealing the dynamics of labeled blood flowing from pial arteries to downstream microvasculature. Resting state CBF peaked in the middle layers which is highly consistent with microvascular density measured from human cortex specimens. Finger tapping induced a robust two-peak laminar profile of CBF increases in the superficial (somatosensory and premotor input) and deep (spinal output) layers of M1, while finger brushing task induced a weaker CBF increase in superficial layers (somatosensory input). We further demonstrated that top-down attention induced a predominant CBF increase in deep layers and a smaller CBF increase on top of the lower baseline CBF in superficial layers of V1 (feedback cortical input), while bottom-up stimulus driven activity peaked in the middle layers (feedforward thalamic input). These quantitative laminar profiles of perfusion activity suggest an important role of M1 superficial layers for the computation of finger movements, and that visual attention may amplify deep layer output to the subcortex.

**Significance Statement:** CBF or microvascular perfusion measured by arterial spin labeling (ASL) is a key parameter for *in vivo* assessment of neurovascular function. Compared to BOLD or VASO fMRI, ASL perfusion contrast offers the unique capability for quantitative CBF measurements both at baseline and during task activation, which is critical for quantitative estimation of metabolic activities tightly related to neuronal activation. We proposed a zoomed 3D ASL technique at 7T for laminar perfusion imaging with high spatial specificity and sensitivity. This technique is able to differentiate and quantify the input/output and feedforward/feedback activities of human motor and visual cortex, thereby providing an important tool for quantitative assessment of neurovascular function and metabolic activities of neural circuits across cortical layers.

## Introduction

It was discovered a century ago that the neocortex in mammals is arranged in six layers that vary across different cortical areas (1, 2). This observation formed the basis for the parcellation of human cortex into separate areas, and further assigned feedforward and feedback pathways to different cortical layers (3, 4). Blood oxygen level dependent (BOLD) fMRI is the most widely used non-invasive imaging modality for studying the dynamics of macroscopic brain networks and mesoscopic brain circuits. It is now feasible to achieve a sub-millimeter spatial resolution using BOLD fMRI at the level of cortical layers at ultrahigh magnetic fields (UHF) of 7T and above (5-7). However, the complex interplay between cerebral blood flow (CBF), cerebral blood volume (CBV) and oxygen metabolism hampers quantitative interpretations of the BOLD signal across cortical layers or depths with different baseline physiology (8). The BOLD signal is also susceptible to contaminations of pial veins on the cortical surface that significantly confounds laminar fMRI. Compared to conventional gradient-echo T2*-weighted BOLD, emerging fMRI methods based on T2-weighted BOLD and CBV contrast have been shown to more precisely map the laminar activities in both animal and human studies (8-13). In particular, vascular space occupancy (VASO) based CBV fMRI at 7T (14, 15) is able to differentiate specific activations in different layers of human motor cortex (M1), as well as the directional functional connectivity of M1 with somatosensory and premotor areas (9). However, VASO only measures relative CBV changes that may be confounded by different baseline CBV values across cortical layers.

CBF or perfusion measured by arterial spin labeling (ASL) is a key parameter for *in vivo* assessment of neurovascular function. The ASL signal is localized close to the site of neural activation as most of the labeled arterial water exchanges with tissue water in capillaries (16). Optical imaging in animals provides evidence that CBF can be precisely and rapidly regulated on an extremely fine scale at the level of capillaries (17-20). Compared to BOLD and CBV fMRI, ASL perfusion contrast offers the unique capability for quantitative CBF measurements both at rest and during task activation, which is critical for quantitative estimation of metabolic activities tightly related to neuronal activation. UHF ASL has the dual benefits of increased signal-to-noise ratio (SNR) that scales with B_0_ field and prolonged tracer half-life (blood T1 = ∼2.1sec at 7T) (21, 22), and therefore may overcome the major limitation of ASL in terms of low SNR. To date, however, the capability for *in vivo* mapping of microvascular perfusion at laminar level remains largely nonexistent, mainly due to the low sensitivity and technical challenges of performing high-resolution ASL at ultrahigh fields.

In this study, we introduced a novel zoomed 3D pseudo-continuous ASL (pCASL) technique at 7T with high spatial resolution (1-mm isotropic) and sensitivity to characterize layer-dependent resting and task activation induced perfusion activity in the human motor and visual cortices. For the first time, multi-delay pCASL was applied to measure variations of arterial transit time (ATT) and resting state CBF across cortical layers, illustrating the dynamics of labeled blood flowing from pial arteries, arterioles to downstream microvasculature in the middle layers of cerebral cortex (23). Zoomed pCASL at the optimal post-labeling delay was then applied on the motor cortex for detecting and quantifying the layer-dependent activity of M1 during finger tapping and finger brushing (SI Fig. S1A). Previous studies have shown that somatosensory and premotor input to M1 largely terminates in the superficial layers (II/III), while cortico-spinal output originates predominantly in the deep layers (Vb/VI) (24-26). However, the precise quantitative sensory input and motor output of M1 remain to be determined. The optimal zoomed pCASL protocol was further applied on the visual cortex to quantify the absolute and relative perfusion changes across cortical layers in response to a visual spatial attention task in V1. While visual attention has been shown to operate through feedback connections along descending visual pathways involving both superficial and deep layers of V1, a quantitative measure is required to evaluate the relative contribution of deep and superficial layers in top-down attention (SI Fig. S1B). In this study, we specifically addressed these unsolved neuroscientific questions using zoomed pCASL perfusion fMRI at 7T.

## Results

### Zoomed pCASL protocol at 7T

Applying pCASL at UHF is challenging due to a high specific absorption ratio (SAR) level of RF power, as well as the B_1_^+^ drop and B_0_ inhomogeneity that affect labeling efficiency (27). We developed an innovative zoomed 3D pCASL technique to achieve a high labeling efficiency and spatial resolution without exceeding the SAR limit. A segmented 3D inner-volume gradient and spin-echo (GRASE) sequence was applied for zoomed pCASL perfusion imaging of a 3D slab of 100×50×24mm^3^ with a high resolution of isotropic 1mm^3^. The flip angels (FA) of GRASE refocusing pulses and scheme for segmented acquisition were optimized along with a single background suppression pulse to minimize spatial blurring, physiological noise and SAR (Fig.1A). The workflow was streamlined by performing maximal-intensity-projection (MIP) of the T1w structural MRI to visualize intracranial arteries, followed by placing the pCASL labeling plane above the circle of Willis (CoW) and simultaneously perpendicular to the M3 segment of middle cerebral artery (MCA), P2 segment of posterior cerebral artery (PCA) and A2 segment of anterior cerebral artery (ACA) (Fig.1B). Compared to conventional ASL labeling plane at carotid arteries, this labeling location has three advantages: 1) B_1_ and B_0_ fields are more homogeneous around the center of the brain; 2) Blood flow velocity is slower, and the pCASL labeling scheme can be optimized to reduce SAR while maintaining sufficient labeling efficiency; 3) The distance between the labeling plane and cortex of interest is reduced resulting in shortened ATT and TR (= 2.8 sec) for perfusion fMRI scans. With this optimized zoomed 3D pCASL, a high labeling efficiency of 82.1% can be achieved with an average SAR of 77.4±8.9% of the first level limit (3.2W/kg on head). Another advantage of the proposed technique is perfusion and T2w BOLD contrasts can be concurrently acquired by pairwise subtraction and summation of label and control images (28).

**Figure 1.**
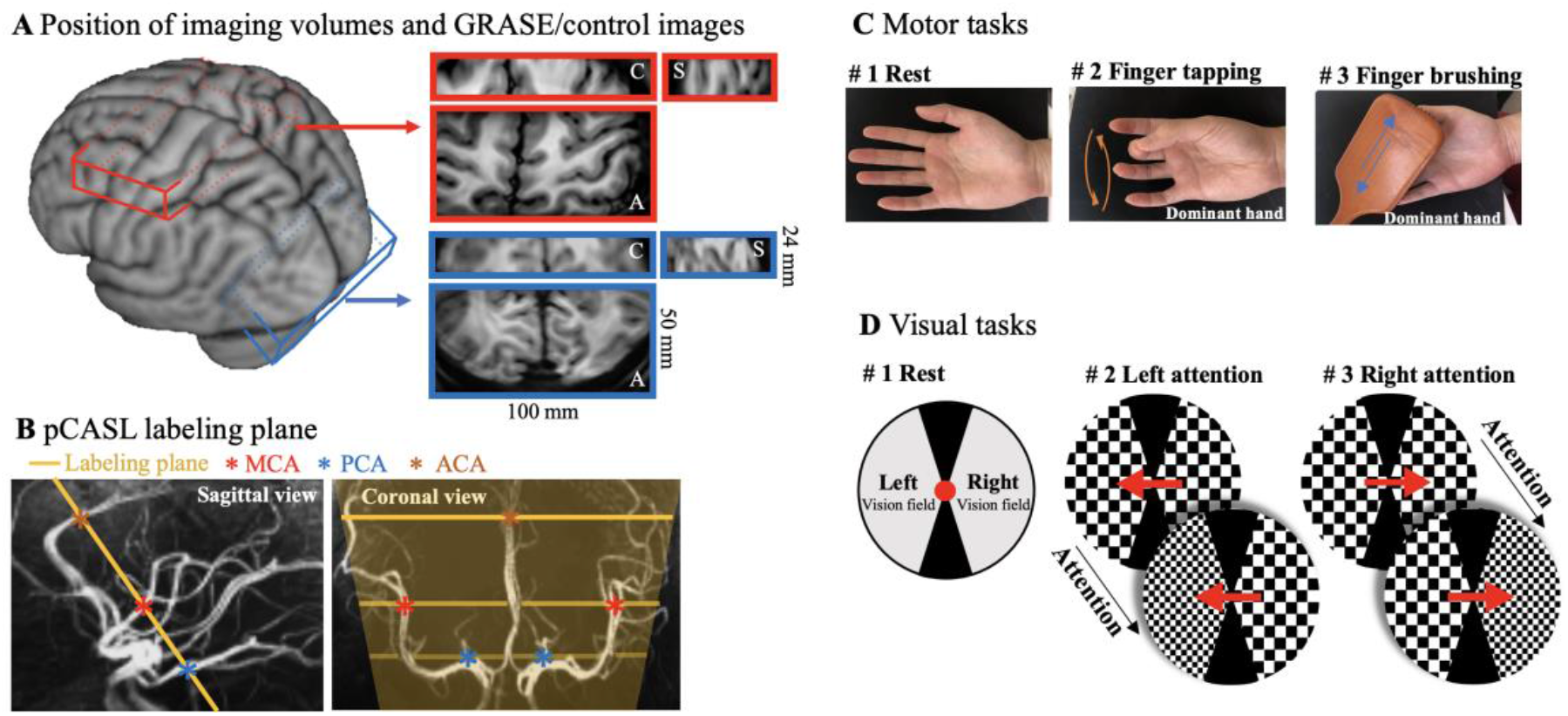
Demonstration of image acquisition and diagrams of functional tasks. **A**. Illustration of imaging volume on a 3D brain surface. A small FOV (100×50×24mm^3^) covering the dominant motor cortex (red) or visual cortex (blue) was acquired with zoomed GRASE. Axial, coronal and sagittal views of GRASE images were shown in enlarged red and blue boxes, respectively. **B**. Illustration of pCASL labeling plane in sagittal (yellow line) and coronal (yellow lines and shade) views. Intracranial arteries were revealed by maximal-intensity-projection (MIP) of the T1w structural MRI and pCASL labeling plane was placed above the circle of Willis (CoW) and simultaneously perpendicular to the M3 segment of middle cerebral artery (MCA), P2 segment of posterior cerebral artery (PCA) and A2 segment of anterior cerebral artery (ACA). **C**. Diagram of motor tasks. Two motor tasks consisted unilateral sequential finger tapping and brushing of the dominant hand (frequency of 2Hz). **D**. Diagram of spatial visual attention tasks. A pair of counter-phase flickering (7.5 Hz, 50% contrast) square wave checkerboards patterns (2 cycles per degree) were presented to the left and the right side of fixation. During the stimulus presentation, the spatial frequency of the two checkerboard patterns changed 7 times randomly and independently. Subjects were asked to pay attention to the cued checkerboard to detect occasional spatial frequency change of the attended stimulus in left or right vision field in two visual task runs.

### Laminar profile of Resting state ATT and CBF

Resting state perfusion scans were performed with the optimized zoomed 3D pCASL at 3 PLDs (500, 1000 and 1500 msec) on the motor and visual cortex, respectively. The ASL imaging volumes were manually positioned to be perpendicular to the ‘omega’ or ‘epsilon’ shaped hand ‘knob’ of M1 (29, 30), and parallel to a flat portion of the calcarine fissure (31) for motor and visual cortex, respectively. Each imaging volume was acquired with 6 segments in 16.8 sec, and 18 pairs of label and control volumes were acquired in 10 min for each PLD. The acquired ASL control images exhibited high contrast between gray and white matter, and demonstrated sufficient coverage for the premotor, supplementary motor area (SMA), S1 and M1 areas of the motor cortex (Fig. 1A & Fig. 2A), as well as for V1 in the calcarine fissure of the visual cortex (Fig. 1A & Fig. 3A), respectively (Perfusion maps of twenty axial slices are shown in SI Fig. S2). Dynamic resting state perfusion maps at three PLDs and calculated ATT and CBF maps of motor and visual cortex are shown in Fig. 2B-D and Fig. 3B-D respectively. The perfusion maps at the short PLD of 500 msec showed heterogeneous signals at the GM/CSF boundary, indicating insufficient time for the labeled blood to flow from the labeling plane to the cerebral cortex, and thus the majority of ASL signal still remained in pial arteries or arterioles. The perfusion maps at the longer PLD of 1000 and 1500 msec showed more homogeneous signal distribution with minimal signal remaining in CSF. As a result, the laminar profiles (Fig. 2E and Fig. 3E) of the perfusion signals at the PLD of 500 msec peaked at the GM/CSF boundary and monotonically decreased towards deep layers, while the peak perfusion signal at the PLD of 1000 and 1500 msec shifted towards middle layers. These findings are consistent with the laminar profiles of ATT (Fig. 2E and Fig. 3E) which were shortest in superficial layers (914.7±12.1 msec in M1 and 912.3±4.4 msec in V1, averaged from four layers at GM/CSF boundary) and increased towards middle to deep layers (1001.4±3.4 msec in M1 and 994.4±0.9 msec in V1, averaged from four layers at GM/WM boundary). Average resting state CBF (corrected for ATT) across cortical depths were 51.9±11.3 ml/100g/min in M1 and 60.3±7.2 ml/100g/min in V1 with peak CBF observed in a middle layer (65.6 ml/100g/min in M1 and 68.5 ml/100g/min in V1, Fig. 2E and Fig. 3E). Peak of resting state CBF is slightly shifted towards middle/deep layers in visual cortex likely due to dense vascular network in granular layer (layer 4) which does not exist in motor cortex. To the best of our knowledge, this is the first time that the dynamics of labeled blood flowing from pial arteries, arterioles to downstream microvasculature is shown *in vivo* on the cerebral cortex. Our results are highly consistent with microvascular density data measured on specimen of human motor and visual cortex (23) (Fig. 2F & Fig. 3F).

**Figure 2.**
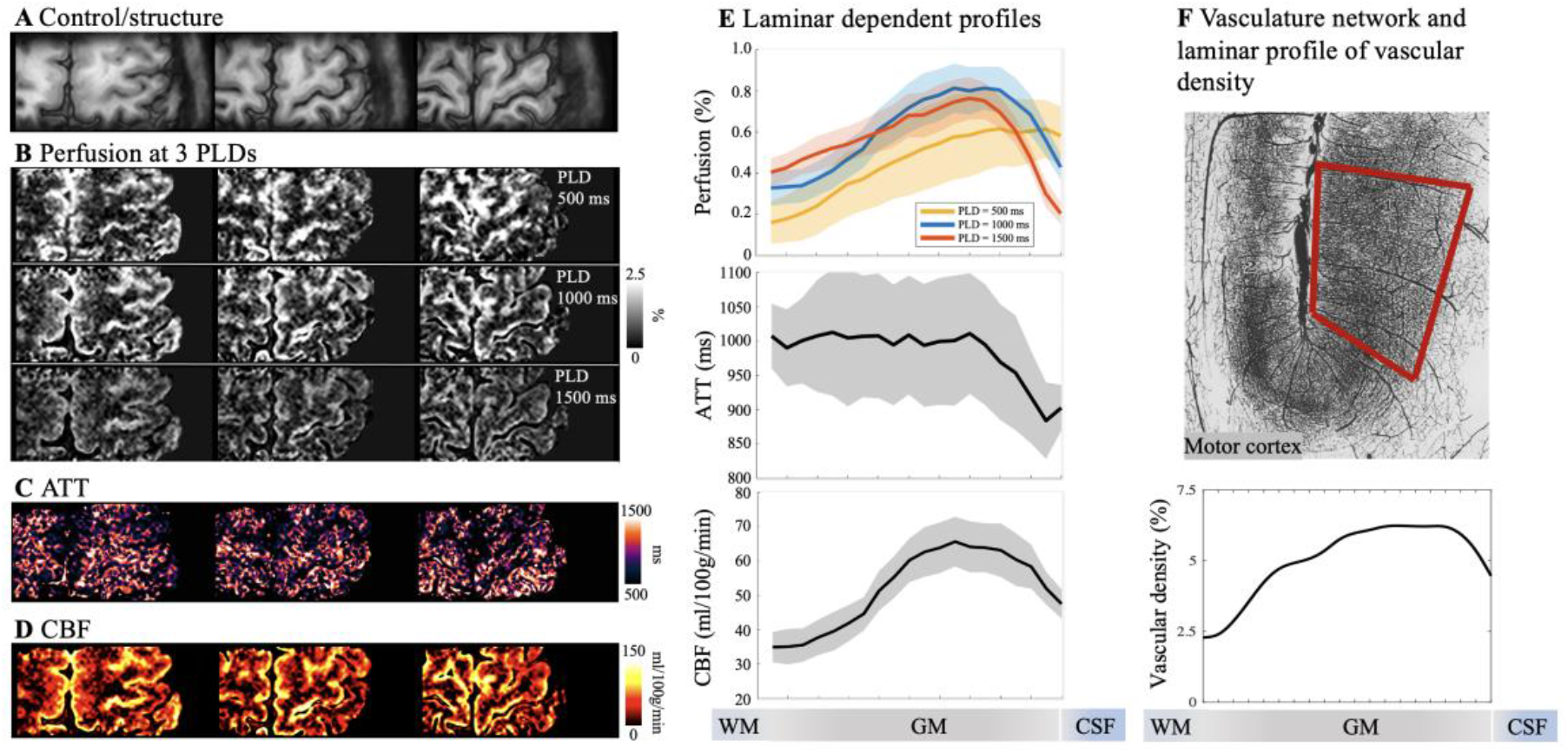
Resting state pCASL perfusion in motor cortex. **A**. Three slices of GRASE/control images demonstrating sufficient coverage for hand knob area in M1. **B**. Resting-state perfusion images acquired at three PLD of 500, 1000 and 1500 msec. Color bar indicates the percentage of perfusion signal in relation to M_0_. Maps of ATT (**C**) and resting state CBF (**D**) were estimated from multi-PLD perfusion signals using a weighted-delay approach. **E**. Laminar profiles of multi-PLD perfusion signals, ATT and resting state CBF. Perfusion signal peaks at GM/CSF boundary at an early PLD of 500 msec, and the perfusion signal peak shifts towards middle layers at later PLDs of 1000 and 1500 msec. ATT was lower at GM/CSF boundary indicating fast transit to pial arteries and became homogeneous in middle and deep layers. CBF was higher in middle layers indicates higher microvascular density. Shaded areas indicate variations across subjects. **F**. Vasculature network obtained from specimen of huma brain in motor cortex (top row, frontal and parietal cortex on right and left side of central sulcus). Vessels are labeled in black and darker area indicate higher vascular density (Figure is adapted from (23)). Laminar profile of relative vascular density (bottom row) calculated from inverted image intensity within the red box.

**Figure 3.**
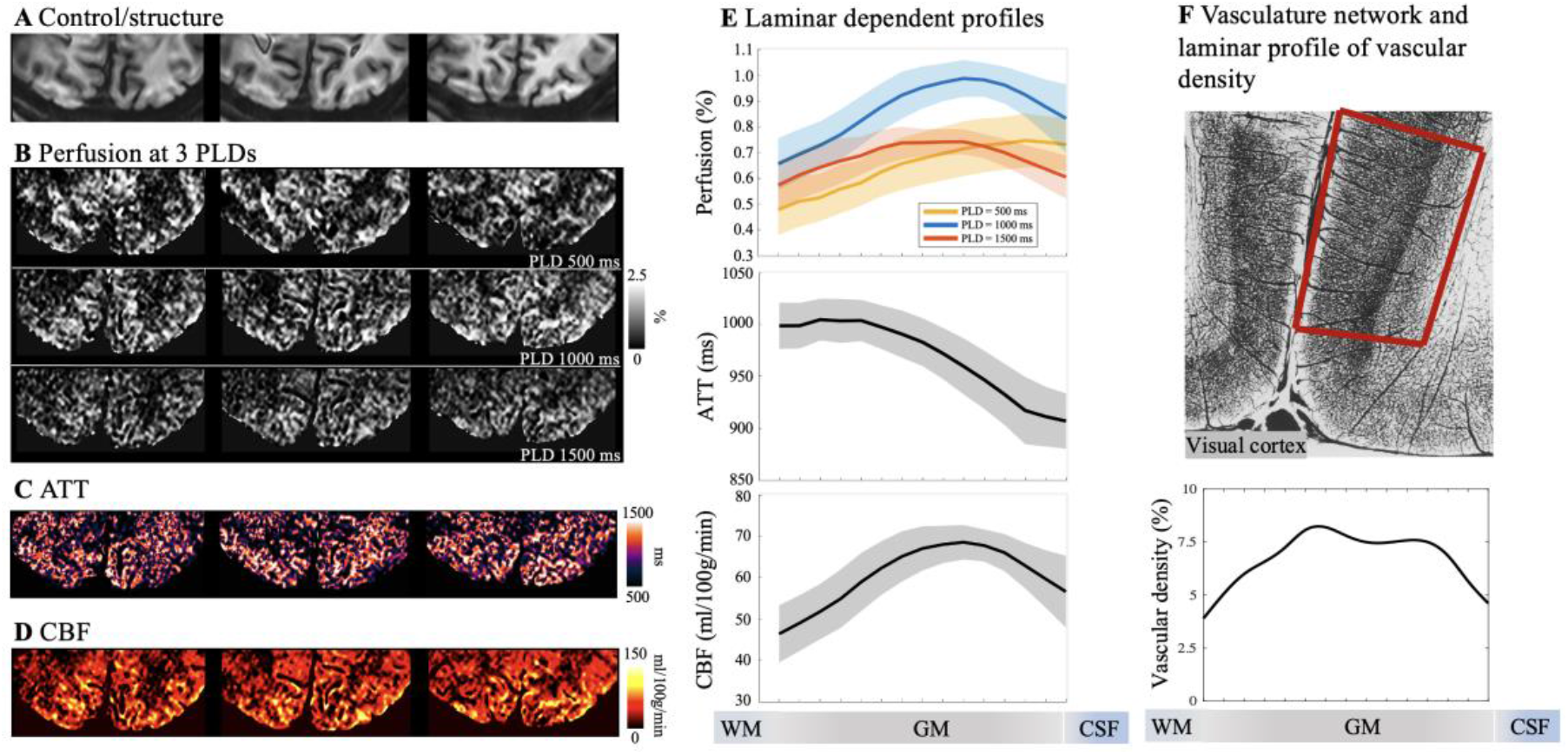
Resting state pCASL perfusion in visual cortex. **A**. Three slices of GRASE/control images demonstrating sufficient coverage for V1. **B**. Resting-state perfusion images acquired at three PLD of 500, 1000 and 1500 msec. Color bar indicates the percentage of perfusion signal in relation to M_0_. Maps of ATT (**C**) and resting state CBF (**D**) were estimated from multi-PLD perfusion signals using a weighted-delay approach. **E**. Laminar profiles of multi-PLD perfusion signals, ATT and resting state CBF. Perfusion signal peaks at GM/CSF boundary at an early PLD of 500 msec, and the perfusion signal peak shifts towards middle layers at later PLDs of 1000 and 1500 msec. ATT was lower at GM/CSF boundary indicating fast transit to pial arteries and became homogeneous in middle and deep layers. CBF was higher in middle layers indicates higher microvascular density. Shaded areas indicate variations across subjects. **F**. Vasculature network obtained from specimen of huma brain in visual/striate cortex (top row). Vessels are labeled in black and darker area indicate higher vascular density (Figure is adapted from (23)). Striate cortex exhibit a distinct characteristic that most dense vascular network is found in the internal granular layer (layer 4). Laminar profile of relative vascular density calculated from inverted image intensity within the red box.

### ASL fMRI for quantifying input and output activity in M1

Perfusion fMRI scans were acquired at the PLD of 1000 ms when the majority of the labeled signal arrives at capillary/tissue space and the dynamic ASL signal reaches its peak (Fig. 2E). Fig. 1C shows the two motor tasks, which consisted of unilateral finger tapping (FT, sequential, frequency of ∼2 Hz) and finger brushing (FB, ∼2Hz) of the dominant hand. CBF maps at resting state and during the two motor tasks are shown in Fig. 4B. The two motor tasks elicited different laminar-dependent activation patterns. Strong and medium CBF increases evoked by the FT and FB tasks can be observed along M1 respectively, as displayed in two zoomed windows (Fig. 4C for absolute CBF change and Fig. 4D for relative CBF change). Both FT induced absolute (ΔCBF) and percentage (%CBF) changes show clearly two peaks in deep (94.8±3.4 ml/100g/min or 159±7.8%) and superficial layers (57.5±1.4 ml/100g/min or 139±3.8%) respectively (Fig. 4E bottom panel, averaged from 5 layers adjacent to peak), consistent with the hypothesis that FT engages neural activity of both somatosensory and premotor input in the superficial layers and motor output in the deep layers (24-26) (SI Fig. S1A). Absolute CBF increase was much larger in the superficial layers than in the deep layers, suggesting that the superficial layers of M1 might involve extensive neural computations integrating the somatosensory and motor planning signals to generate motor output in the deep layers. Interestingly, this finding was not observed in CBV-weighted VASO fMRI of human M1 which showed comparable activation in the superficial and deep layers during FT (24). Statistical significance of the two-peak pattern was validated by R^2^ _diff_ which measures the likelihood that a profile consists of two isolated distributions instead of one. This double peak pattern is highly reliable (P = 0.011 for ΔCBF and P < 0.0002 for %CBF, SI Fig. S8) and can be observed in all participants (see individual results in SI appendix, Fig. S3). FB-induced CBF increase was much smaller averaged cross cortical layers compared to that of the FT task. The perfusion response mainly peaked in superficial layers (Fig. 4E bottom panel, 38.8±2.7 ml/100g/min or 69.3±5.7% and 15.3±0.9 ml/100g/min or 45.2±1.5% in the superficial and deep layers (averaged from 5 layers adjacent to peak) respectively), consistent with the hypothesis that FB primarily engages somatosensory input and minimal motor output (SI Fig. S1A). These results demonstrate the high spatial specificity of ASL, capable of resolving and further quantifying layer-dependent input and output activity in human M1. Both FT and FB induced BOLD signal changes are mostly in superficial layers (Fig. 4E top right). FT induced BOLD response shows a weak increase in deep layers; however, the two-peak pattern was not significant (P = 0.47) due to the large BOLD signal change in superficial layers. Furthermore, correction of BOLD signal which was on the order of 1-2% had minimal effect on perfusion signal changes (Fig. 4E lower right dashed trace). Our results are highly consistent with CBV fMRI data, nevertheless ASL fMRI provides additional measurements of both absolute and relative CBF changes with FT and FB inducing 73.6±12.5 ml/100g/min and 26.0±6.4 ml/100g/min or 160.6±28.6% and 51.5±22.0% CBF increase across all cortical depths. This is the first time that such detailed quantitative information on both absolute and relative perfusion activation related to the input and output activity of the motor cortex is visualized in vivo.

**Figure 4.**
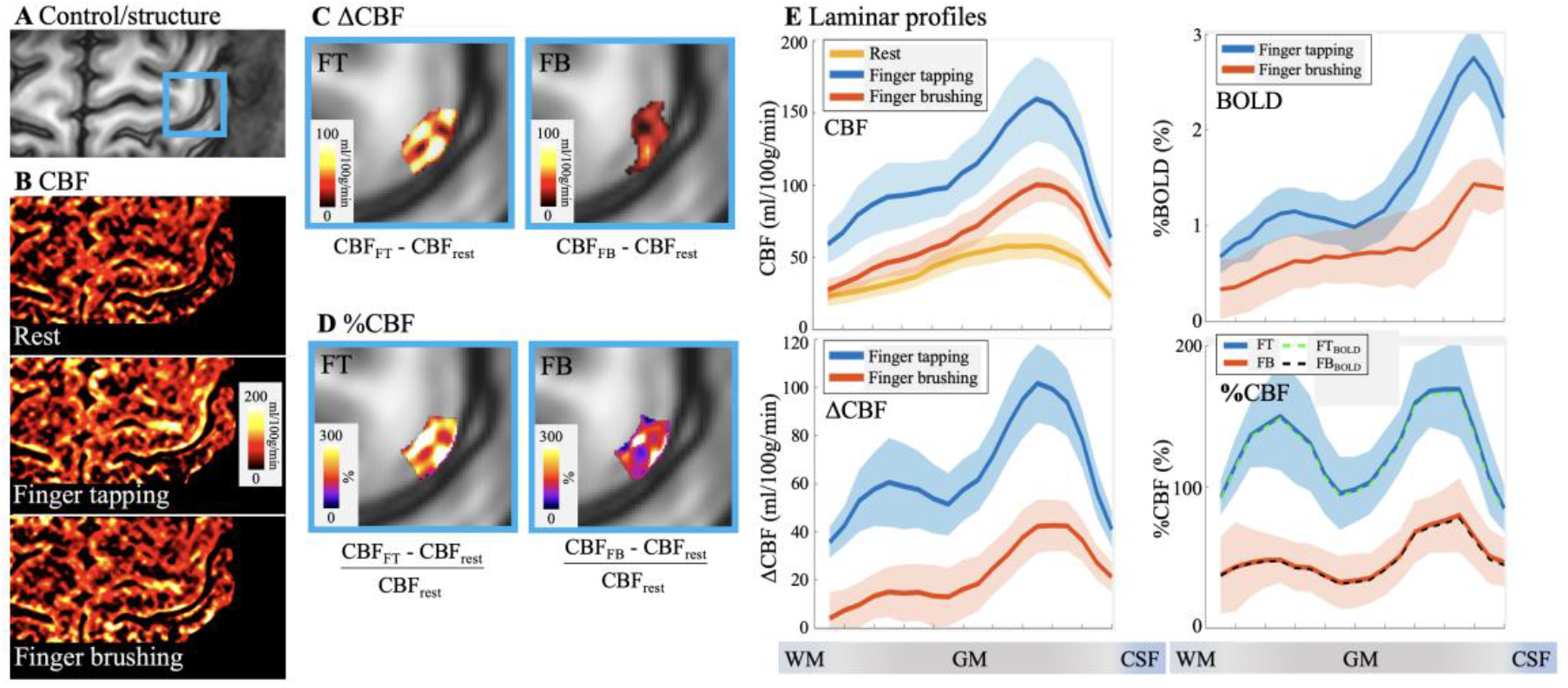
Motor tasks activated pCASL perfusion. **A**. GRASE image with hand knob area highlighted in blue box. **B**. CBF maps at rest (top), finger tapping and finger brushing. Absolute (**C**) and percentage (**D**) CBF increase evoked by the FT and FB tasks are highlighted in M1 hand knob area (blue boxes). **E**. Laminar profile of CBF at rest and two motor tasks (top left), percentage BOLD changes (top right), absolute (bottom left) and percentage (bottom right, dashed line (FT_BOLD_ and FB_BOLD_) was adjusted for BOLD effects) CBF changes induced by two motor tasks. Both FT and FB induced BOLD signal changes are mostly in superficial layers. FT induced BOLD response shows a weak increase in deep layers; however, the two-peak pattern was not significant (P = 0.47) due to the large BOLD signal change in superficial layers. FT induced CBF increase shows a ‘two-peak’ pattern (P = 0.011 and P < 0.0002 for significance test of two-peak pattern of ΔCBF and %CBF, respectively) which corresponds to sensory input (superficial) and motor output (deep) layers respectively. FB induced CBF response shows a weaker increase in superficial layers which corresponds to exteroception sensory input and minimal motor output. Shaded areas indicate variations across subjects

### ASL fMRI for quantifying feedforward and feedback activity in V1

We further applied 3D zoomed pCASL for perfusion fMRI with a visual spatial attention task to resolve and quantify the feedforward and feedback activities of human V1. As shown in Fig. 1D of the task paradigm, subjects were required to pay attention to the cued checkerboard pattern presented on either left or right visual field to detect occasional spatial frequency change of the attended stimulus (32). Resting state CBF and visual attention activated CBF maps are shown in Fig. 5B. The attended and unattended visual stimuli induced strong CBF increases localized along the gyri and sulci of visual cortex. Absolute CBF increases to unattended visual stimuli over baseline and attended over unattended stimuli are displayed in two zoomed windows (Fig. 5C). Fig. 5D shows the relative CBF change evoked by unattended visual stimuli over baseline (left column) and the multiplicative effect of attention modulation map (right column) defined as the ratio between CBF increase evoked by top-down attention (attended - unattended) and stimulus-driven response (unattended - baseline). Fig. 6E shows the laminar profile of CBF at rest and with unattended and attended visual stimuli (top left), BOLD changes (top right), absolute and relative CBF changes induced by unattended visual stimuli (middle panel, BOLD corrected relative CBF changes are also shown), absolute CBF changes induced by top-down attention (bottom left), and CBF and BOLD response to attention modulation (bottom right). Both absolute and relative CBF increase evoked by unattended visual stimuli peak in middle layers, consistent with the hypothesis that that feedforward visual input from the lateral geniculate of the thalamus mainly reaches the middle layers of V1 (33, 34) (SI Fig. S1B). Absolute CBF increase evoked by top-down attention shows a large peak in deep layers while the CBF profile of multiplicative attention modulation shows a two-peak pattern (P = 0.039 of two-peak pattern significance test, SI Fig. S8) in both deep (19.0±1.7%, two-layer average) and superficial layers (14.5±1.8%, two-layer average) versus middle layers (8.3±0.5%, two-layer average). These results are consistent with the hypothesis that attention operates through feedback pathways involving both deep and superficial layers (32, 33) (SI Fig. S1B). The single-peak pattern of unattended visual stimulation and double peak pattern of attention modulation is highly reliable and was observed in all participants (see individual results in SI appendix, Fig. S4). In contrast, BOLD profile of attention modulation is relatively flat across layers with a slight increase towards GM/CSF boundary (P = 0.063 of two-peak pattern significance test).

**Figure 5.**
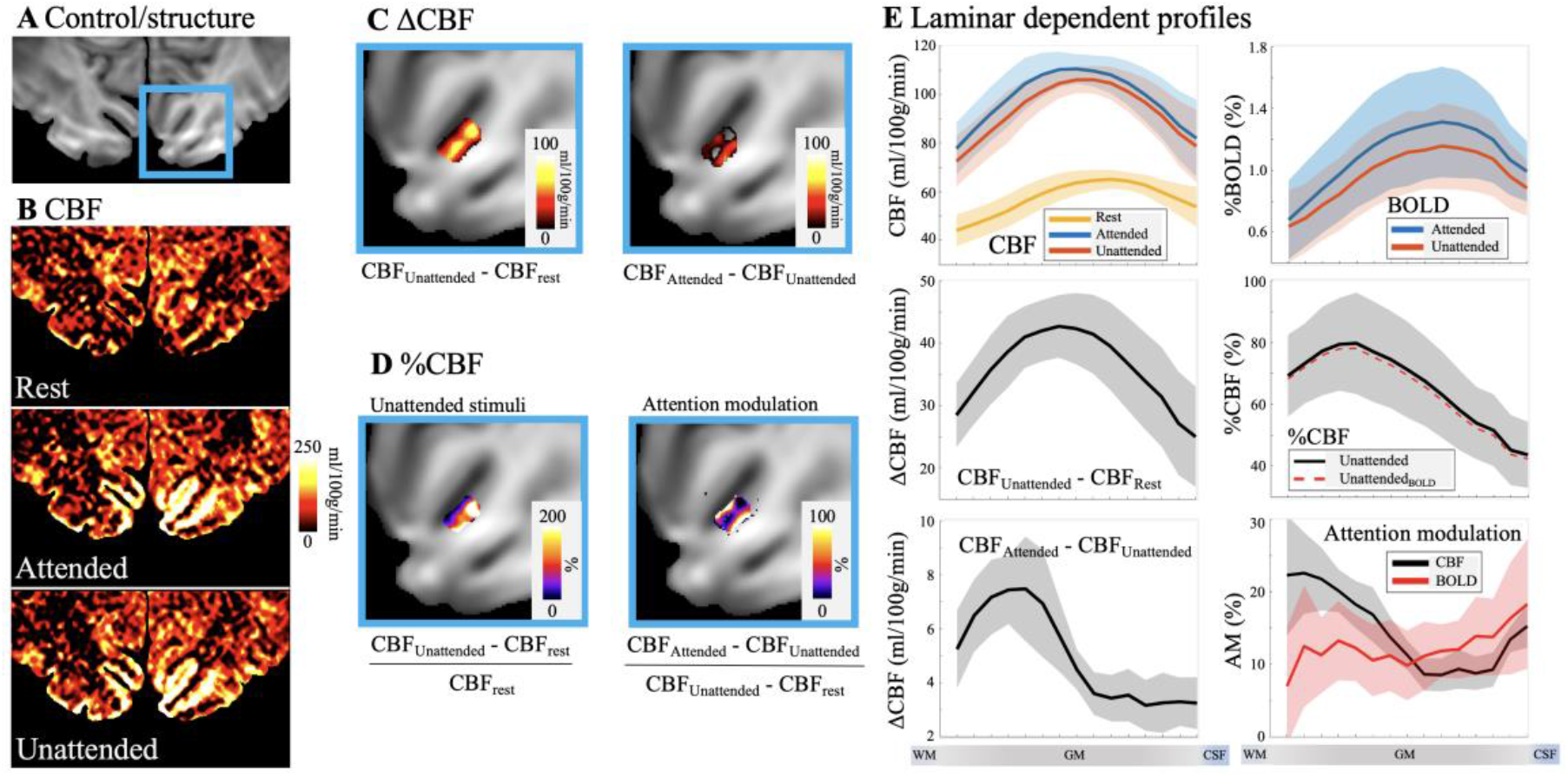
pCASL perfusion at spatial attention tasks. **A**. GRASE image with V1 highlighted in blue box. **B**. CBF maps at rest (top), attended and unattended visual stimuli. **C**. Absolute CBF increase evoked by unattended visual stimuli (left column) and spatial attention (right column) in V1. **D**. Relative CBF increase evoked by unattended visual stimuli (left column) and attention modulation (right column). **E**. Laminar profile of CBF at rest and with unattended and attended visual stimuli (top left), percentage BOLD changes (top right), absolute and relative CBF changes induced by unattended visual stimuli (middle panel, dashed lines were adjusted for BOLD effects), absolute CBF changes induced by spatial attention (bottom left) and CBF and BOLD response to attention modulation (bottom right). Both absolute and relative CBF increase evoked by unattended visual stimuli peak in middle layers. Absolute CBF increase evoked by spatial attention shows a large peak in deep layers while CBF profile of attention modulation shows a two-peak patten (P = 0.039 of two-peak pattern significance test). BOLD profile of attention modulation is relatively homogeneous across layers with a slight increase towards GM/CSF boundary (P = 0.063 of two-peak pattern significance test). Shaded areas indicate variations across subjects.

## Discussion

We demonstrated that multi-delay ASL can reveal perfusion dynamics and quantify ATT and resting state CBF across cortical layers. Vascular architecture in human brain is specialized to maintain and regulate blood supply through surface pial arterioles (35), the subsurface microvascular bed and penetrating arterioles bridging the surface and subsurface networks (36). We found baseline ATT was shortest at GM and CSF boundaries indicating fast arrival of blood at pial arteries on cortical surfaces. ATT in middle and deep layers were rather homogeneous indicating a fast transit through penetrating arterioles to the subsurface microvascular bed (23, 36, 37). The laminar profile of baseline CBF exhibited a single peak around the middle layers and monotonically decreased towards deep and superficial layers in both M1 and V1. The CBF profile matches well with the highest capillary density observed in the middle layers as reported in anatomic studies in specimens of human brain tissue (23, 38). Additionally, we observed the peak of resting state CBF in visual cortex is more into deep layers as compared to motor cortex, which likely corresponds to a granular layer with high vascular density in visual cortex. Pathological ATT or CBF heterogeneity across cortical layers might be associated with disturbances in endoneurial metabolism or capillary morphology and implicate a number of diseases including stroke (37, 39), ischemia (40), Alzheimer’s disease (41), traumatic brain injury (42), and diabetic nephropathy (43). Studying laminar dependent perfusion dynamics may provide meaningful insights into the progression of neurological diseases.

We further demonstrated the sensitivity and spatial specificity of the proposed ASL fMRI in differentiating and quantifying input and output-driven neural activities in M1. Compared to the concurrently acquired T2w BOLD contrast, ASL fMRI was able to detect two separate peaks in superficial and deep layers of M1 (Fig. 4) across all participants (SI Fig. S3), while the two-peak pattern was not significant in BOLD signals. This result is highly consistent with VASO fMRI data (14), supporting that both CBF and CBV based laminar fMRI is able to differentiate input and output activities of human M1. Nevertheless, CBF-based fMRI provides both absolute and relative perfusion activation, thereby more quantitative information on the magnitude of neural activation across cortical layers. In contrast, VASO only measures relative CBV changes that may be confounded by different baseline CBV values across cortical layers. Furthermore, the proposed ASL fMRI is robust to potential BOLD contamination which is on the order of 1-2%. The quantitative measurement of CBF increases to the finger tapping task revealed a larger peak in the superficial layers than in the deep layers, suggesting an important role of M1 superficial layers integrating the input signals from somatosensory and premotor cortex to generate motor output in the deep layers.

ASL fMRI was also applied with a visual spatial attention task to detect and quantify feedforward and feedback activities in the visual cortex. We observed multiplicative attention modulation in both deep and superficial layers while unattended visual stimuli induced peak CBF increase mainly in the middle layers (Fig. 5). Furthermore, the quantitative perfusion changes induced by top-down attention shows a single large peak in the deep layers. The peak of attention modulation in the deep layers was not clearly visible in concurrently acquired T2w BOLD contrast. Our result is highly consistent with the known neuroanatomy of feedforward and feedback visual pathways (27), further supporting the high sensitivity and spatial specificity of CBF-based fMRI. The superficial and deep layers of V1 receive feedback inputs from higher cortical areas. The superficial layers also send output to higher order visual cortex, while the deep layers send output to the subcortical visual areas including the LGN, pulvinar and SC, etc (34, 44). The multiplicative effect of attention modulation in both superficial and deep layers suggests that the control of attention operates thorough descending feedback pathway from higher cortical areas, while the predominant increase of absolute CBF in the deep layers suggest that the effect of attention mainly amplifies output signals to the subcortex (44) (SI Fig. S1B).

Recent studies demonstrated the feasibility of performing ASL fMRI with high spatial resolutions (45, 46). At ultra-high fields, ASL has the dual benefits of long tracer half-life determined by blood T1 (47) and a super-linear relationship between SNR and field strength (48), resulting in a ∼3 fold SNR increase at 7T as compared to 3T. However, shorter arterial blood T2 (68 msec) at 7T (49) places a strict limit on the choice of TE to minimize signal loss as well as blurring along slice direction. We achieved minimal TE and echo-train length using zoomed imaging with half FOV along phase encoding direction, partial Fourier acquisition and segmented readout. Physiological noises in dynamic perfusion signals were minimized by a principle component analysis (PCA)-based denoising algorithm (50) and signal blurring was further minimized by POCS reconstruction and a SVD based deblurring algorithm. A diagram of the deblurring process is shown in SI Fig. S5. The resultant point-spread function (PSF) of our method was 1.37 pixel which is sharper than or comparable to the PSFs of BOLD and CBV fMRI (51).

Both imaging coverage and temporal resolution of the proposed technique can be further improved by utilizing accelerated image acquisition and constrained reconstructions (52, 53). It is also feasible to perform perfusion based functional connectivity analysis of brain network dynamics with increased coverage and temporal resolution (54). In this study we observed strong perfusion based functional connectivity between M1 and associated brain regions including premotor, SMA and S1 (see individual functional connectivity map in SI Fig. S6). The proposed technique can be further extended to obtain concurrent CBF, BOLD and CBV measurements for mapping metabolic activities (55). With the advent of even higher field strength of 10.5T MRI machine (56) and high density array coils (57), the sensitivity and resolution of zoomed pCASL can be further improved.

In conclusion, we demonstrated high spatial specificity and sensitivity of laminar perfusion fMRI using 3D zoomed pCASL at 7T in detecting and quantifying the input versus output and feedforward versus feedback activities of neural circuits in the motor and visual cortices. With the unique capability for quantitative CBF measurements both at baseline and during task activation, high-resolution ASL perfusion fMRI at 7T provides an important tool for *in vivo* assessment of neurovascular function and metabolic activities of neural circuits across cortical layers.

## Materials and Methods

### Human participants

Six right-handed participants (age = 28.2±3.5 years, 2 males and 4 females) underwent motor task experiments and six participants with normal or corrected to normal vision (age = 27.7±3.7 years, 3 males and 3 females) underwent spatial visual attention experiments, respectively. All subjects refrained from caffeine intake three hours before the scan. All participants provided written informed consents according to a protocol approved by the Institutional Review Board (IRB) of the University of Southern California.

### Experimental protocol and session setup

Experiments were conducted on a 7 Tesla Terra scanner (Siemens Healthineers, Erlangen, Germany), using a single-channel transmit and 32-channel receive head coil (Nova Medical, Wilmington, MA, USA). Third order shimming was performed before ASL scans to improve B_0_ field homogeneity. Head motion was minimized by placing cushions on top and two sides of head and taping participant’s chin to coil. Experienced participants were recruited in this study and overall framewise displacement of ASL scans was less than 0.2 mm (SI Fig. S7).

Imaging parameters for ASL scans were: FOV=100×50 mm^2^, 24 slices (33% oversampling, 6/8 partial Fourier), 1-mm isotropic resolution, 2 and 3 segments along phase and partition directions, TE=26.78 msec, echo train length=214.2 msec, labeling duration=1280 msec, TR=2800 msec. FAs of 8 refocusing pulses were 120^0^ and the first FA was increased to 150^0^ for signal stabilization. Zoomed imaging was achieved by switching excitation and refocusing gradients between slice and phase directions for inner-volume acquisition (Fig.1A). One non-selective HS pulses was used for suppressing background signal. M1 and V1 were identified from uniform (UNI) MP2RAGE (0.7-mm isotropic resolution, inversion time = 1000/3200 ms, scan time=9 min 46 sec) images. ASL imaging volumes were placed perpendicular to the ‘omega’ or ‘epsilon’ shaped hand ‘knob’ of M1 (29, 30) and parallel to a flat portion calcarine fissure (31) for motor and visual task experiments, respectively.

Intracranial vasculature was revealed by maximal-intensity-projection (MIP) of the UNI MP2RAGE images, and pCASL labeling plane was placed above the circle of Willis (CoW) and simultaneously perpendicular to the M3 segment of MCA, P2 segment of PCA and A2 segment of ACA (Fig.1B). Flow velocity at the labeled vessels was measured by ECG-gated phase-contrast MRI in two subjects. Average blood flow velocity at labeled arteries was 18.4±4.0 cm/sec. Average B1 at labeling was 74.0±11.5% of ideal B1.

Motor and visual task experiments were conducted in two separate sessions. Each session included MP2RAGE, resting state ASL scans at two PLDs (500/1500 msec, 8 min 24 sec) and four runs of motor or visual task ASL scans (10 min 4 sec) at PLD=1000 msec. Total scan time was less than 70 mins per session. Each task run consisted of nine interleaved blocks (67.2 sec per block) with counterbalanced order of tasks across participants.

Fig. 1C shows the paradigm of motor tasks, which consisted of unilateral finger tapping (sequential, frequency of 2 Hz) and finger brushing of the dominant hand. Participants were instructed to remain relaxed with their hands facing up during the brushing task. Participants’ fingers were brushed back and forth by a custom-made MRI safe brush at a frequency of 2 Hz. Fig. 1D shows the stimuli and paradigm of the visual task. Visual stimuli were generated in MATLAB (Mathworks Inc.) with psychophysics toolbox (https://www.psychtoolbox.net/). Stimuli were presented with the BOLDscreen 32 LCD (Cambridge Research Systems Ltd, Rochester, UK) installed at the end of MRI bore. Participants viewed the stimuli through a mirror mounted on top of the head coil. Subjects were required to keep fixation during the experiment. Before stimulus presentation, a central cue was presented at fixation for 1 second, then a pair of counter-phase flickering (7.5 Hz, 50% contrast) square wave checkerboards patterns (2 cycles per degree) were presented to the left and the right side of fixation. The size of the checkerboard discs were 7 degrees in diameter, presented at an eccentricity of 6 degrees. During the stimulus presentation, the spatial frequency of the two checkerboard patterns changed 7 times randomly and independently. Subjects were asked to pay attention to the cued checkerboard to detect occasional spatial frequency change of the attended stimulus in left or right vision field in two task runs. Successful detection of the cued pattern within 2 secs was recorded by pressing a button box. Average accuracy was 73.7±13.9% across participants.

### Signal processing

Partial Fourier reconstruction was done by the Projection Onto Convex Sets (POCS) method (58). Dynamic ASL image volumes from resting state and task ASL runs were realigned and co-registered using SPM12 (Functional Imaging Laboratory, University College London, UK). Spatial weightings were applied for M1 and V1 to avoid distortion or imperfect slice profile in FOV boundaries. Average framewise displacement (FD) of resting state and functional ASL runs were 0.08±0.03 and 0.19±0.07 mm in motor cortex and 0.09±0.03 and 0.13±0.03 in visual cortex (Parameters of rigid head motion are shown in SI Fig. S7). A boundary based algorithm was used to co-register ASL image volumes and structural MP2RAGE image (59). Control and label images were pairwise subtracted and summed to obtain perfusion and T2w BOLD images respectively (28). Dynamic signal fluctuations induced by motion and physiological noises were minimized by a PCA-based algorithm (50). Signal decay in k-space was prominent in partition direction due to long echo train length. k-space profile along partition direction was estimated by the extended phase graph (EPG) method (60) with inputs of TE, flip angle train and T2 of arterial blood (68 msec (49)). The point spread function (PSF) of perfusion signal blurring along slice direction was calculated by 1-D Fourier transform of the k-space signal. A singular value decomposition (SVD) based deblurring algorithm was implemented to reduce FWHM of PSF from 2.15 mm to 1.37 mm (SI Fig. S5).

Resting state (PLD=1000 msec) and task activation perfusion signals were extracted and re-combined from four task ASL runs. ATT and resting state CBF map were simultaneously estimated from resting state perfusion signal at three PLDs using a weighted-delay approach (61). Task activation CBF was calculated according to (62) incorporating ATT. CBF maps with visual attention stimuli in left or right visual fields were calculated, and corresponding hemispheres with and without attention were separated and re-combined to produce attended and unattended CBF maps.

### Segmentation of cortical layers in M1 and V1

ASL images were upscaled to a finer grid of 0.25×0.25×0.5 mm^3^ resolution to avoid singularities at the edges in angular voxel space. The borderlines of CSF/GM and GM/WM in M1 hand-knob area were manually drawn on ASL control images. Twenty cortical layers of M1 were segmented by LAYNII (63) using the equi-volume layering approach (64). Segmentation of cortical layers in V1 was done on reconstructed surface. MP2RAGE volume was skull removed and segmented into WM, GM and CSF using FreeSurfer (7.1.1). AFNI/SUMA and custom python/matlab codes were used to generate the equi-volume surfaces (https://github.com/herrlich10/mripy) between WM and pial surfaces. Fifteen cortical layers in V1 were then projected back to volume space for analysis. The equi-volume layering approach used in this study has been shown to have higher accuracy than the equi-distance model by taking into account the local curvature of pial and white matter surfaces into account (64). Considering the difference between cortical thickness in M1 (∼4mm) and V1 (∼2-3 mm) (65), fifteen and twenty layers were segmented in V1 and M1 respectively. With a nominal 1 mm resolution, the effective resolution allows it to detect only ∼4 independent data points. Hence, the defined cortical depths do not represent the MRI effective resolution.

### Significance test for two-peak activation pattern

Two-peak patterns were observed in CBF FT activation and attention modulation profiles. To test the significance of the two-peak pattern, we calculated R^2^_diff_ for each profile. R^2^_diff_ was defined as the difference between the adjust R^2^ estimated from curve fitting assuming two or single Gaussian distributions. Larger R^2^_diff_ indicates the profile can be better described as two peaks instead of a single peak. Statistical significance of the R^2^_diff_ was estimated by 5000 Boot strapping runs with random noise in each cortical layer estimated by inter-subject variation. P value was calculated as the probability that R^2^_diff_ score of profile can be explained by noise only (SI Fig. S8). P < 0.05 was considered as significant. Although other measures including Hartigan’s dip statistic (HDS), bimodality coefficient (BC), Akaike’s information (66) and Larkin’s-F-scores (24, 67) were proposed for multimodality versus unimodality test, the proposed R^2^_diff_ score was well-defined for double-peak detection and more robust to misclassification.

## Acknowledgments

This work was supported by the National Institute of Health (NIH) grant UH3-NS100614, S10-OD025312, R01-NS114382 and R01-EB028297.

