## Supplementary material for "Laminar perfusion imaging with zoomed arterial spin labeling at 7 Tesla": SI Fig. S1

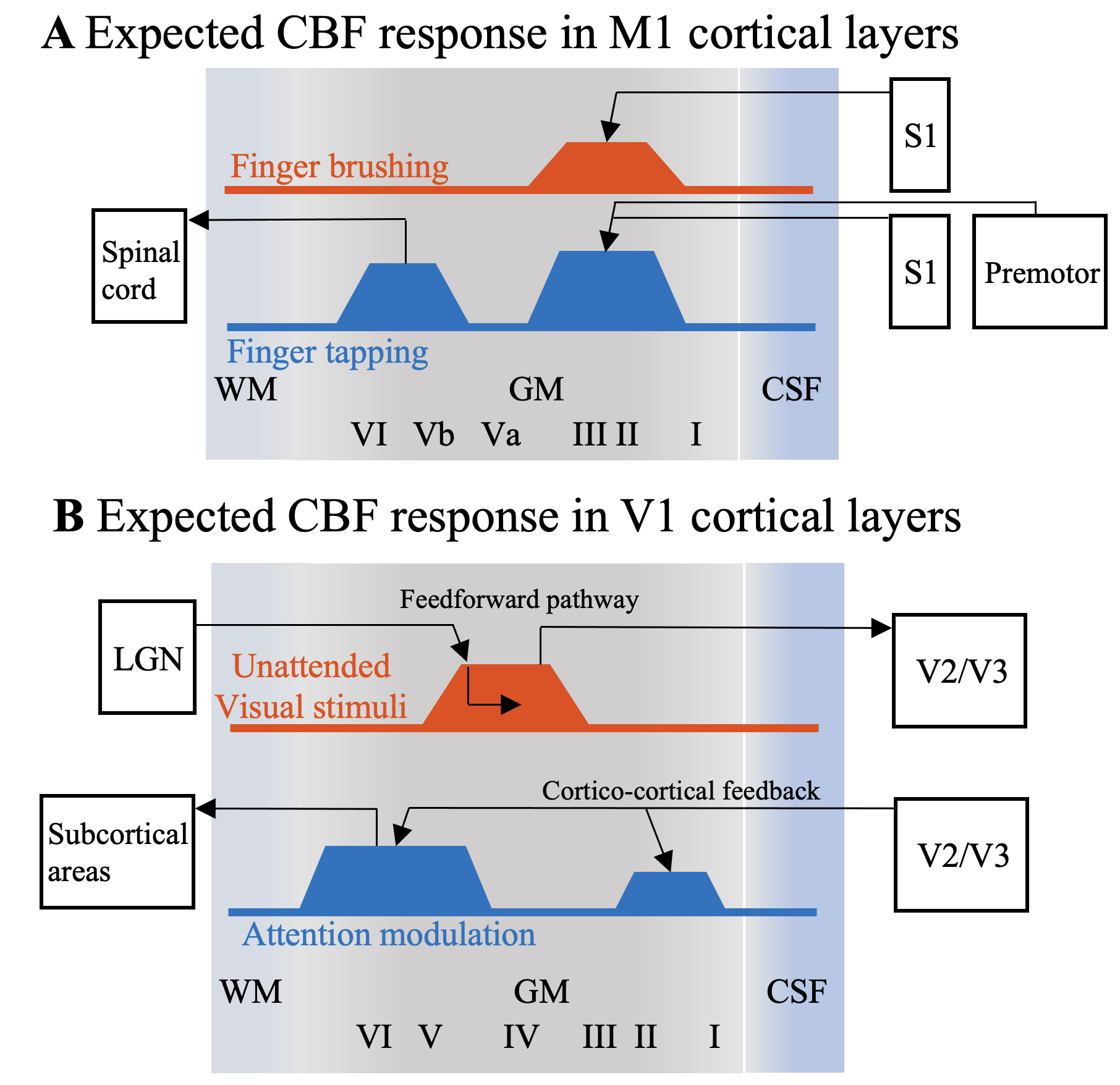

**Figure S1. A.** Demonstration of the expected CBF increase evoked by neuronal activity in response to input and output stimulus. FT induces neuronal activity in both superficial layers (both proprioception and exteroception sensory input from S1 and premotor cortex) and deep layers (motor output), while FB induces weaker neuronal activity in superficial layers (exteroception S1 input) with minimal motor output. (Diagram adapted from (1)). **B.** Demonstration of the expected CBF increase evoked by neuronal activity in response to unattended visual stimuli and attention modulation. Unattended visual stimulation primarily involves feedforward activity in middle layers of the visual cortex, while attention involves descending feedback pathway from higher cortical areas and amplifies output signals to the subcortical areas (Diagram adapted from (2, 3)). Abbreviations: LGN, lateral geniculate nucleus.
