## Supplementary material for "Laminar perfusion imaging with zoomed arterial spin labeling at 7 Tesla": SI Fig. S2

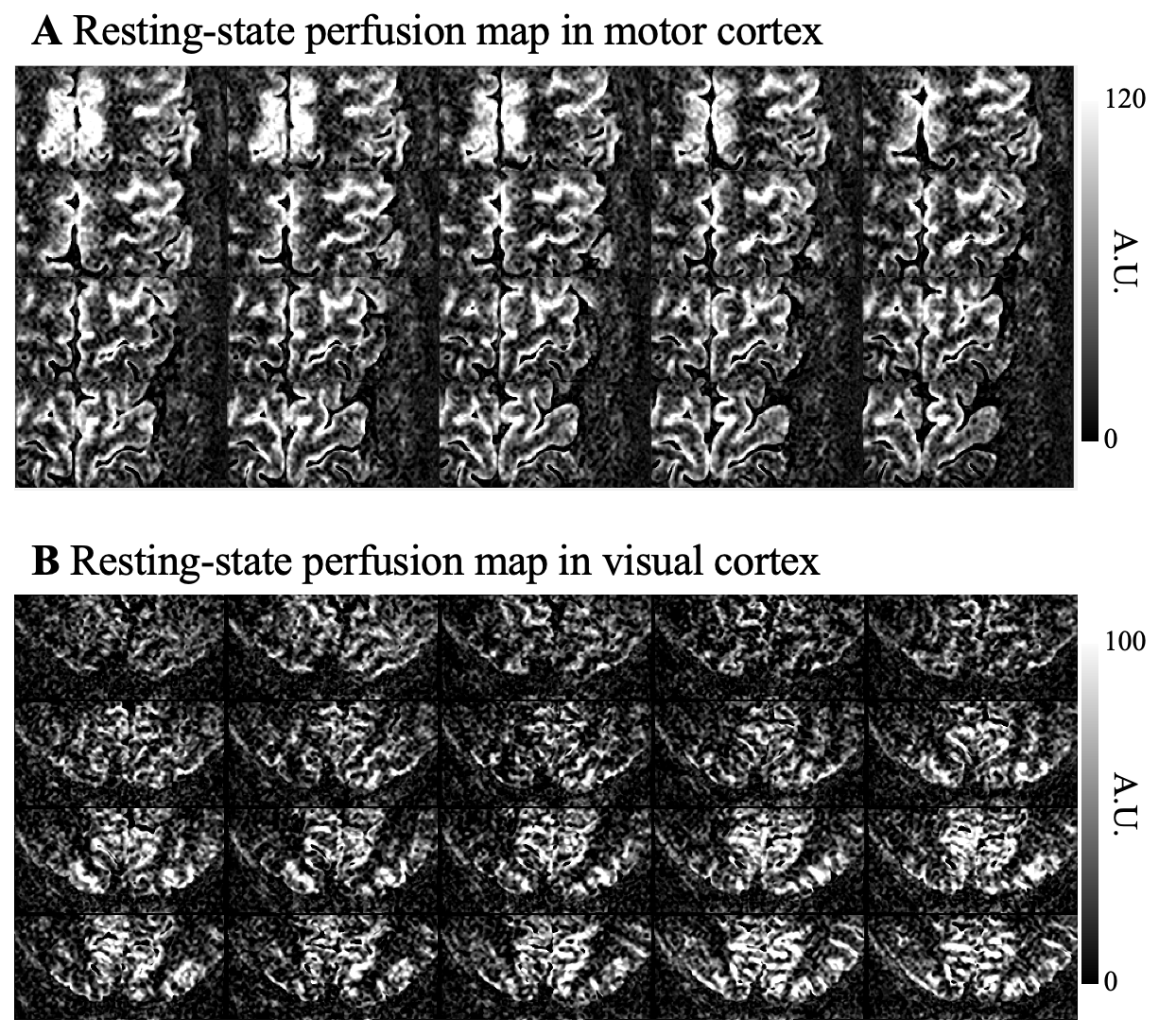

**Figure S2.** Center twenty slices of resting state perfusion maps acquired at PLD of 1000 msec in motor cortex (**A**) and visual cortex (**B**).
