## Supplementary material for "Laminar perfusion imaging with zoomed arterial spin labeling at 7 Tesla": SI Fig. S3

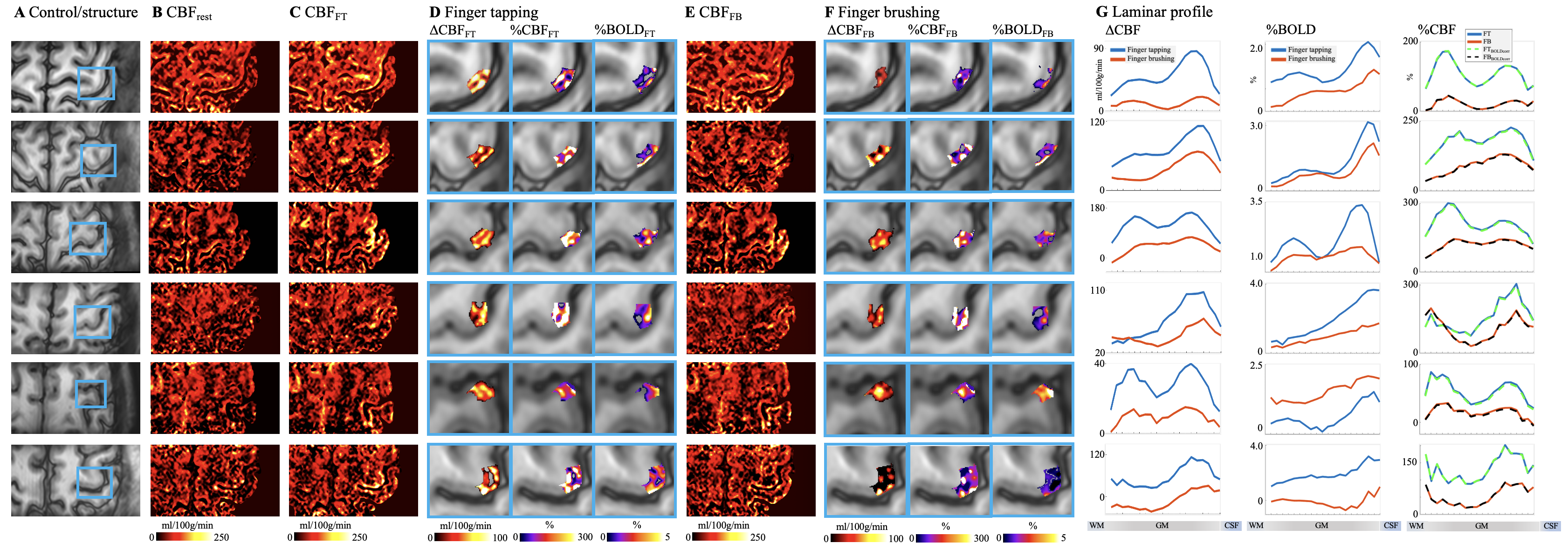

**Figure S3.** Individual results (displayed in each row) of ASL perfusion with two motor tasks. **A.** Control images with M1 hand-knob areas highlighted in blue boxes. Resting stage CBF and CBF maps at FT and FB tasks are shown in **B**, **C** and **E**. Absolute CBF, relative CBF and BOLD signal change in M1 induced by FT and FB tasks are shown **D** and **F**. Laminar profiles of Absolute CBF (left column), BOLD signal (middle column) and relative CBF change (right column, dashed traces were corrected for BOLD effect) evoked by FT (blue trace) and FB (orange trace) are shown in **G**.
