## Supplementary material for "Laminar perfusion imaging with zoomed arterial spin labeling at 7 Tesla": SI Fig. S4

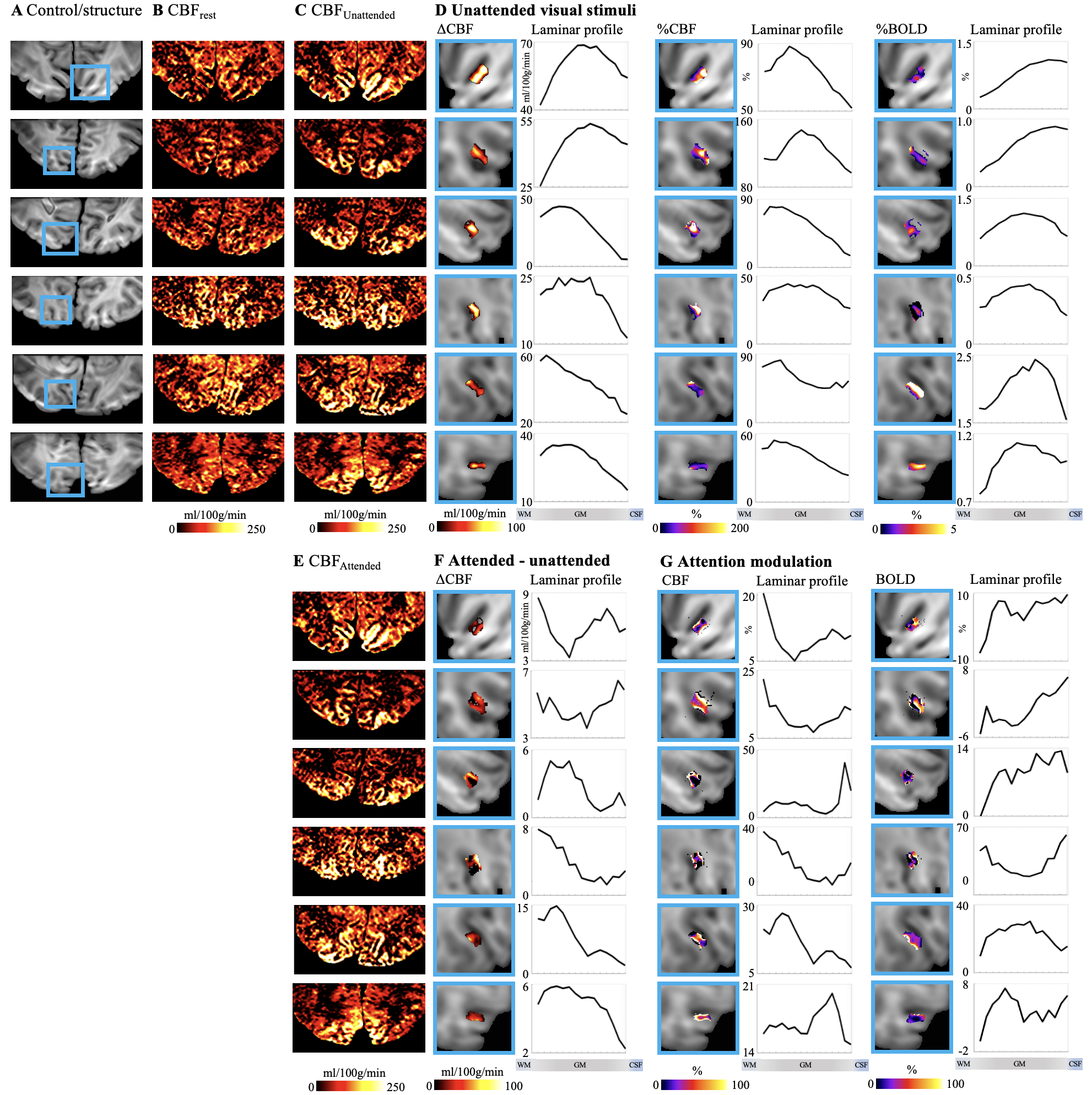

**Figure S4.** Individual results (displayed in each row) of ASL perfusion with spatial visual attention tasks. **A.** Control images with V1 highlighted in blue boxes. Resting stage CBF and CBF maps with unattended and attended visual stimuli are shown in **B**, **C** and **E**, respectively. Maps and laminar profiles of absolute CBF, relative CBF and BOLD signal change induced by unattended visual stimuli are shown in **D**. Maps and laminar profiles of absolute CBF change induced by attention are shown in **F**. Maps and laminar profiles of CBF and BOLD signal change corresponding to attention modulation are shown in **G**. CBF or BOLD signal changes are highlighted in blue box for illustration purpose. Laminar profiles were calculated from larger ROIs covering entire V1.
