## Supplementary material for "Laminar perfusion imaging with zoomed arterial spin labeling at 7 Tesla": SI Fig. S5

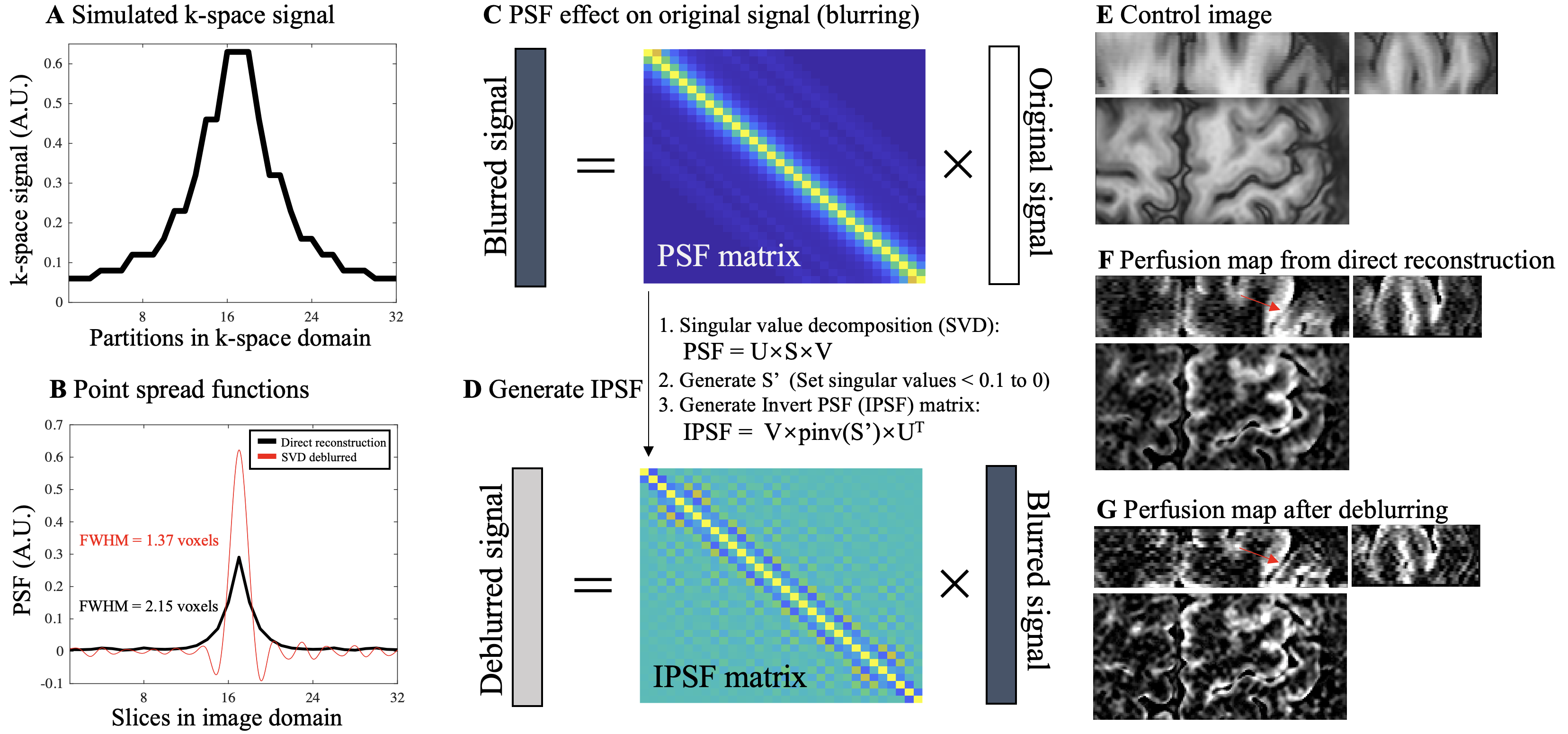

**Figure S5.** Diagram of a singular value decomposition (SVD) based deblurring process for ASL perfusion signal. **A.** k-space signal along 32 (24+33%oversampling) partitions was simulated by the extended phase graph (EPG) method with inputs of TE (26.78 msec), GRASE flip angle train and T2 of arterial blood (68 msec). Due to signal decay towards k-space boundaries, blurring of perfusion signal happens along slice direction. **B.** PSF, which indicates the degree of spatial blurring, of the perfusion signal was calculated by 1-D Fourier transform of the k-space signal and shown as the black trace (FWHM = 2.15 voxels). Red trace is the PSF after the deblurring process, as explained in **C** and **D** (FWHM = 1.37 voxels). **C.** Acquired perfusion signal (vector with size of 32×1) along slice direction can be expressed as the convolution between original signal (no blurring, vector with size of 32×1) and 1-D PSF. Convolution can be reformulated as multiplication between a PSF matrix (size of 32×32) and the original signal vector. **D.** Deblurring can be considered as an inverse problem and deblurred signal can be obtained by multiplying an inverse PSF (IPSF) matrix and blurred signal. We utilized a singular value decomposition (SVD) based approach to solve IPSF. Smaller singular values (elements of S < 0.1) were discarded to minimize noise amplification. Deblurred PSF (red trace in **B**) can be calculated by multiplying IPSF and PSF from direction reconstruction (black trace in **B**). **E**, **F** and **G** show coronal (top left), sagittal (top right) and axial (bottom) views of control images, perfusion map from direct reconstruction (blurred along Z direction) and perfusion map processed by the proposed deblurring approach, respectively. Higher contrast between central sulcus and perfusion signal in M1 and S1 can be observed in the deblurred perfusion map (indicated by red arrows).
