## Supplementary material for "Laminar perfusion imaging with zoomed arterial spin labeling at 7 Tesla": SI Fig. S6

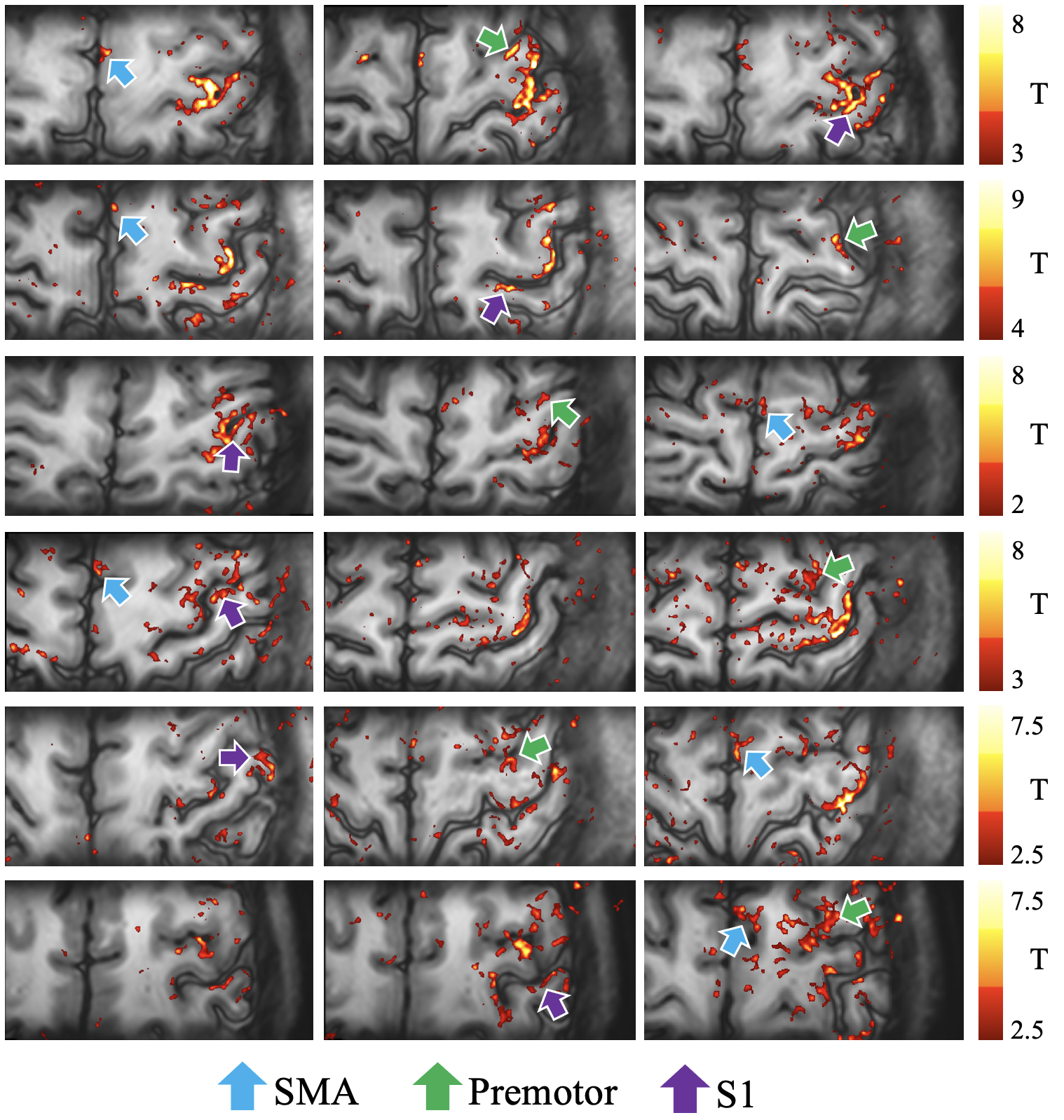

**Figure S6.** Perfusion based functional connectivity maps across six participants (each row). Dynamic perfusion signals from all tasks were concatenated and fitted to the signal fluctuations obtained from a seed region M1 (SPM12). In all participants M1 show strong functional connectivity between SMA (blue arrows), promotor area (green arrows), S1 (purple). The scale bar indicates the T value of regression between voxel-wise fluctuations and seed signals. T > 2.38, 3.21 and 3.90 indicate P < 0.01, 0.001 and 0.0001, respectively. Abbreviations, SMA, supplementary motor area, M1, primary motor cortex, S1, primary sensory cortex.
