## Supplementary material for "Laminar perfusion imaging with zoomed arterial spin labeling at 7 Tesla": SI Fig. S7

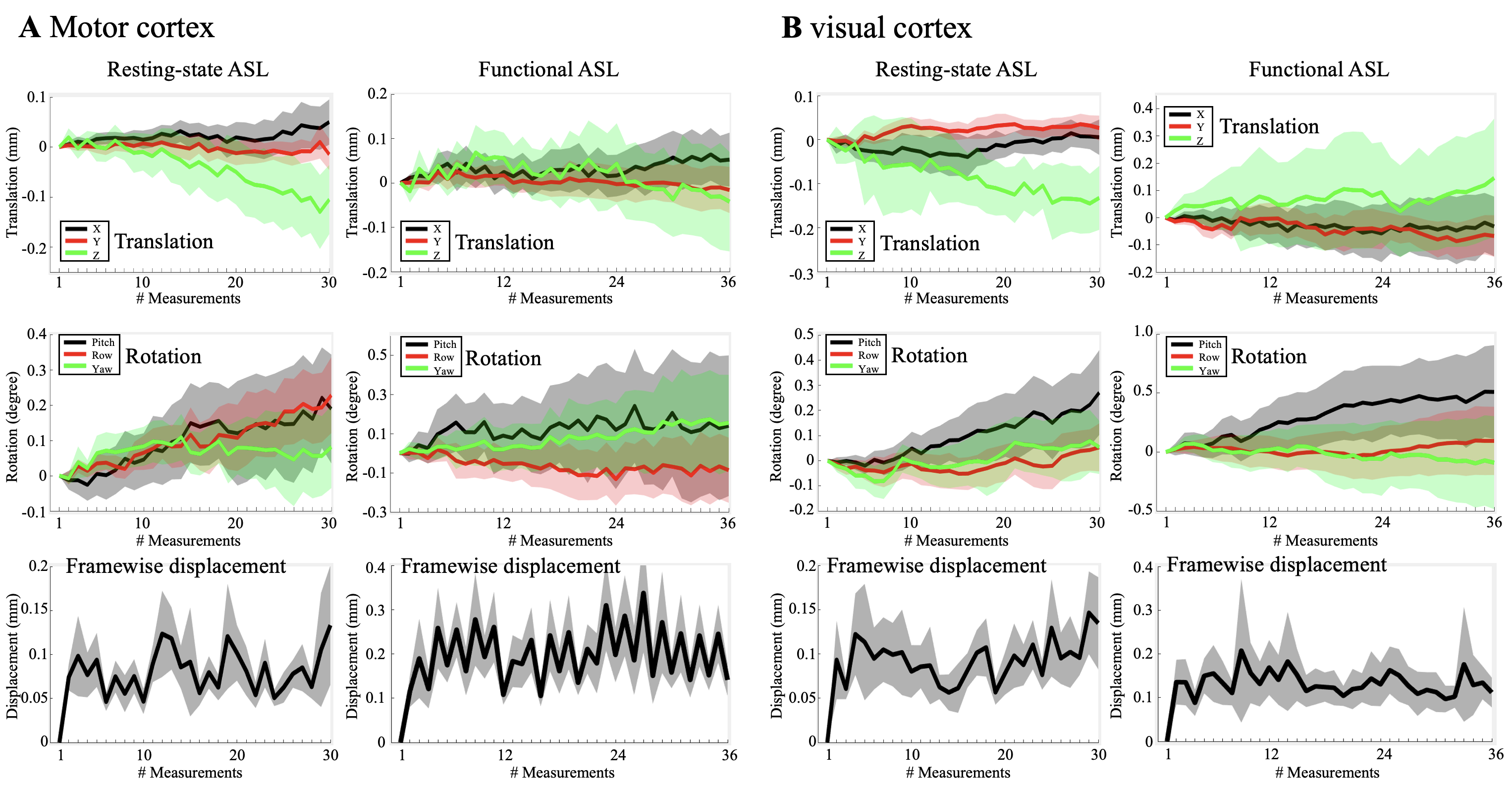

**Figure S7.** Plots of estimated six-degree rigid head motion parameters during two resting state and four functional ASL scans in motor cortex (**A**) and visual cortex (**B**). Translation (mm) and rotation (degree) are shown in the first and second rows, respectively. Framewise displacement (mm) was calculated as the displacement of one farthermost voxel on the volume (50/25/12 mm away from the center in X/Y/Z directions) given the translation and rotation parameters. Shaded areas indicate variations between individual ASL runs.
