## Supplementary material for "Laminar perfusion imaging with zoomed arterial spin labeling at 7 Tesla": SI Fig. S8

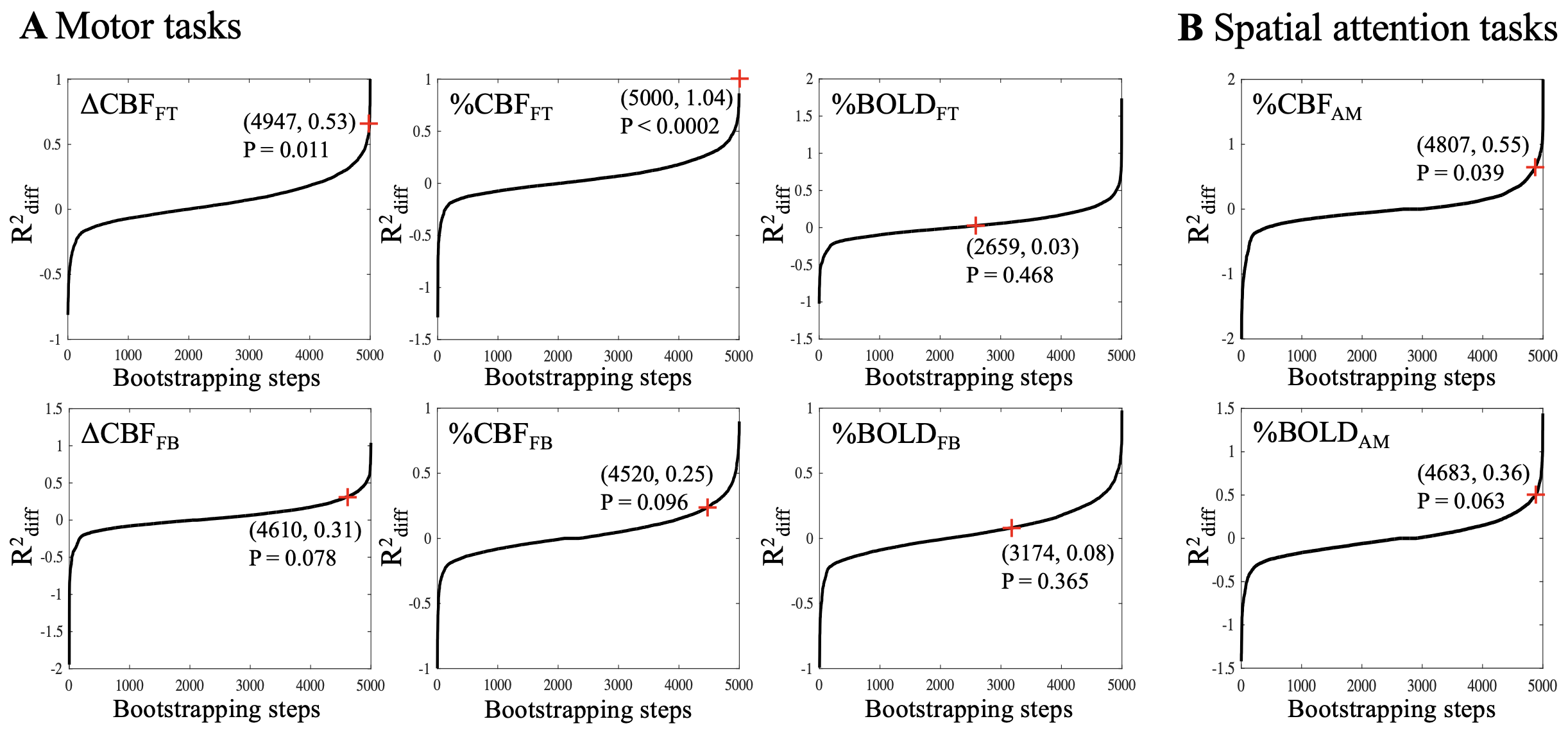

**Figure S8.** Plots of 5000 Boot strapping samples and the R^2^_diff_ statistic of the noise estimated from inter-subject variations in motor cortex (**A**) and visual cortex (**B**). Larger R^2^_diff_ indicates higher likelihood that the profile consists two distributions instead of one. Red cross indicates the rank of the profile R^2^_diff_ score estimated from FT and FB induced absolute CBF change (first column), relative CBF change (second column), BOLD signal change (third column) and CBF/BOLD profiles of AM (fourth column). Number of noise R^2^_diff_ scores smaller than profile R^2^_diff_ score and profile R^2^_diff_ score are listed in the parentheses. P value was calculated as the probability that noise R^2^_diff_ scores greater than the profile R^2^_diff_ score.
